# A unified framework for quantitative splicing and transcript prediction

**DOI:** 10.64898/2026.10.08.757673

**Authors:** Kishore Jaganathan, Jianbin Chen, Xin Liu, Yan Zhang, Yujin Kim, Abhimanyu Banerjee, Petko P. Fiziev, Naomi Wilcox, Ibrahim I. Taskiran, Yichao Cai, Emilie M. Wigdor, Guo-Liang Chew, Grace Png, Jeremy F. McRae, Aniketh Janardhan Reddy, Kuan-Hao Chao, Rami Mehio, Stephan J. Sanders, Kyle Kai-How Farh

## Abstract

RNA splicing is frequently disrupted by genetic variation, contributing to a wide range of human diseases. Identifying the variants responsible for altered splicing is therefore crucial but has long been challenging. Deep learning models that predict splicing from genomic sequence have substantially advanced their detection, but accurately resolving their quantitative effects on splicing and the resulting transcripts remains an unmet need. Here we introduce SpliceAI2, a deep learning model trained end-to-end on splice site, splice junction, and transcript usage across multiple mammalian species. SpliceAI2 reconstructs complete transcripts directly from genomic sequence and considerably improves identification of splice-altering variants and quantification of their effects in diverse benchmarks spanning cryptic splicing, splicing quantitative trait loci, and massively parallel reporter assays. Human population datasets provide independent validation of these predicted functional effects through evidence of natural selection and associated changes in plasma protein abundance. Application to a rare disease cohort reveals that predicted splice-altering variants are highly enriched in disease-associated genes and account for 15% of the excess genetic burden, with approximately half residing in deep intronic regions likely missed by exome sequencing. Together, these findings establish SpliceAI2 as a major step forward in interpreting splice-altering variants across the human genome.

## Main

RNA splicing generates much of the transcript diversity observed in humans through a complex splicing code^1,2^. Genetic variants often disrupt this code, resulting in aberrant splicing that compromises normal biological function. Aberrant splicing is increasingly recognized as a widespread mechanism contributing to common and rare diseases^3–5^. Accurately identifying splice-altering variants is essential for understanding the genetic basis of disease and realizing the full diagnostic potential of whole-genome sequencing^6,7^.

Transcriptome-wide characterization of splicing using RNA sequencing (RNA-seq) has enabled deep learning models to learn key features of the splicing code directly from genomic sequence. SpliceAI established sequence-based splice site prediction as a powerful framework for interpreting splice-altering variants and has been incorporated into guidelines for the clinical interpretation of non-coding variants^8,9^. Pangolin and DeltaSplice subsequently expanded this framework to quantitative prediction of splice site usage^10,11^, and more recently Splam and AlphaGenome broadened it to include splice junction usage^12,13^. Despite these advances, quantitative prediction of variant effects on splicing remains limited, with complete transcript outcomes difficult to resolve.

### Hierarchical model of RNA splicing

We introduce SpliceAI2, a deep learning model that processes approximately 200 kb of genomic sequence to predict position-wise splice site usage and pairwise splice junction usage, and integrates these predictions to infer complete transcripts and their usage (Fig. 1a and Extended Data Fig. 1). We trained SpliceAI2 end-to-end on splice site and splice junction usage from 314,745 RNA-seq samples spanning human and nine additional mammalian species, together with transcript usage from 330 long-read RNA-seq samples in human and mouse^14,15^.

**Fig. 1.**
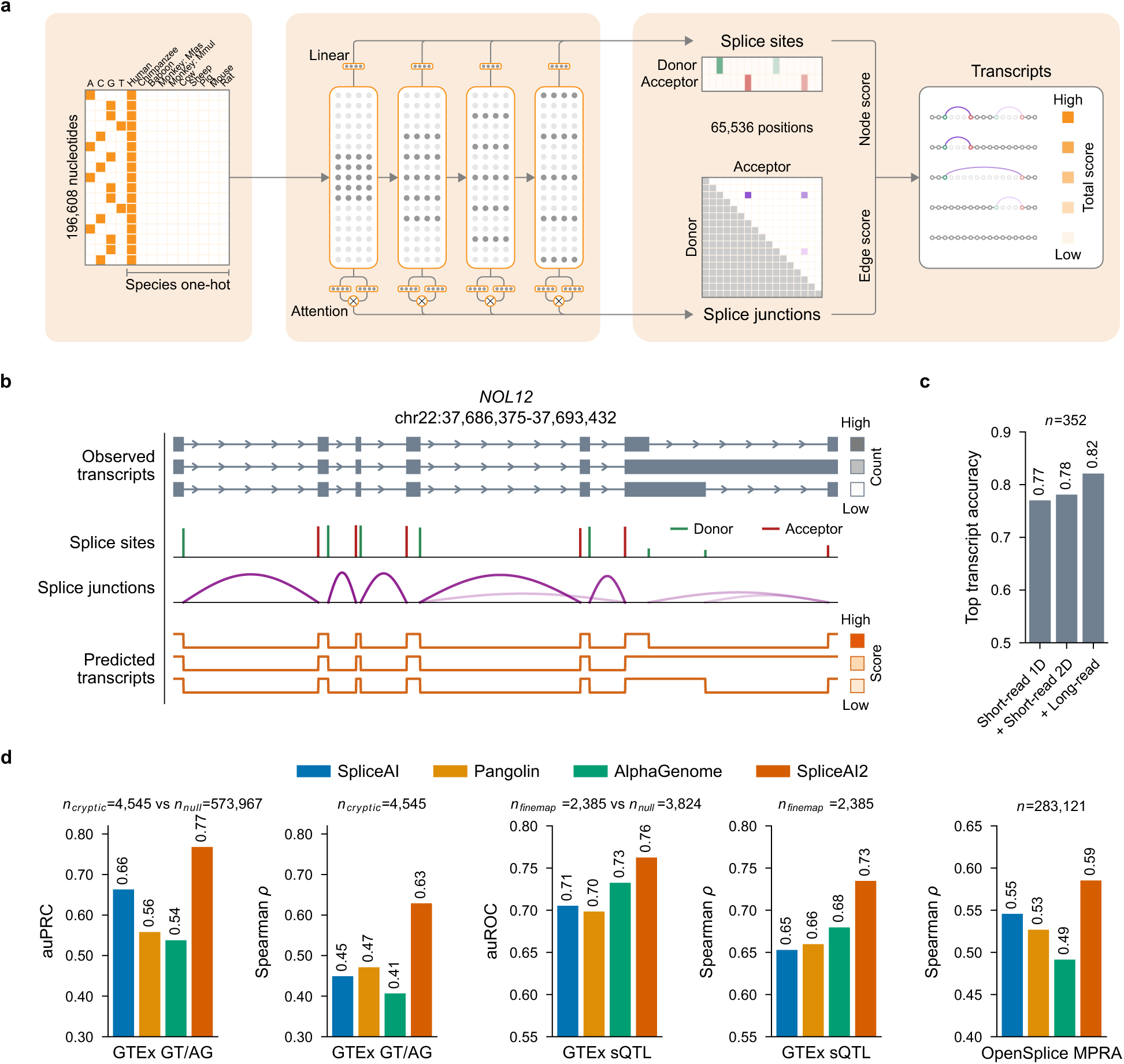
Hierarchical prediction of splice site, splice junction, and transcript usage. **a,** Schematic of the SpliceAI2 architecture, which processes 196,608 bp of genomic sequence and species identity to predict splice site usage across the central 65,536 bp and pairwise splice junction usage among the 1,024 highest scoring splice site positions. These predictions are integrated into a splice graph from which complete transcripts and their usage are inferred using dynamic programming. **b,** Representative example showing observed transcripts and corresponding SpliceAI2 predictions for the held-out *NOL12* gene, with exact reconstruction and correct usage ranking of the three most abundant transcripts. **c,** Percentage of held-out genes with perfect reconstruction of the most abundant transcript, showing that training with long-read transcript usage data through dynamic programming improved performance. Models used for **b** and **c** were trained exclusively on human data because, unlike variant effect evaluations, reference sequence evaluations are susceptible to cross-species leakage through orthologous genes. **d,** Comparison of SpliceAI2 with SpliceAI, Pangolin, and AlphaGenome across three independent variant effect benchmarks spanning GTEx cryptic splice site detection and usage prediction, sQTL classification and effect size prediction, and OpenSplice MPRA effect size prediction.

To integrate splice site and splice junction predictions at the transcript level, SpliceAI2 constructs a splice graph in which candidate splice sites are represented as nodes and candidate splice junctions as edges, with node and edge weights determined by their respective predicted usage. Each candidate transcript corresponds to a path through this graph, with its predicted usage determined by the sum of the node and edge weights along the path. During training, predicted usage is optimized using observed usage across all candidate transcripts, with dynamic programming enabling efficient computation over this exponentially large set^16–21^. SpliceAI2 perfectly reconstructed the most abundant transcript for 82% of held-out genes, compared with 78% when trained without using long-read RNA-seq data (Fig. 1b,c). These findings indicate that short-read data provide an effective foundation for transcript reconstruction, with long-read data further improving reconstruction accuracy.

We compared SpliceAI2 with SpliceAI, Pangolin, and AlphaGenome across three independent splice variant effect benchmarks, excluding essential splice variants throughout (Fig. 1d). We first identified all rare variants creating a GT or AG dinucleotide in the Genotype-Tissue Expression (GTEx) whole-genome sequencing (WGS) data and used matched RNA-seq to identify those that activated cryptic splice sites and quantify the resulting splice site usage^22^. The resulting benchmark comprised 4,545 cryptic splice variants and 573,967 null variants, providing a comprehensive test of cryptic splice site detection in a genome-wide setting. SpliceAI2 considerably outperformed the next best model, SpliceAI (auPRC, 0.77 versus 0.66), with the largest improvements observed for deep intronic variants (Extended Data Fig. 2). Among the 4,545 cryptic splice variants, SpliceAI2 also quantified splice site usage considerably more accurately than the next best model, Pangolin (Spearman *ρ*, 0.63 versus 0.47). We next used GTEx splicing quantitative trait loci (sQTLs) to evaluate whether models could distinguish variants prioritized by fine-mapping from those not prioritized. SpliceAI2 outperformed all competing models on this task (auROC, 0.76 versus 0.73 for AlphaGenome). Among the fine-mapped variants, SpliceAI2 predictions correlated more strongly with observed sQTL effect sizes than predictions from other models (Spearman *ρ*, 0.73 versus 0.68 for AlphaGenome).

Finally, we evaluated the models on the recently published OpenSplice massively parallel reporter assay (MPRA) benchmark^23^, where SpliceAI2 showed the strongest agreement with experimentally measured variant effects (Spearman *ρ*, 0.59 versus 0.55 for SpliceAI).

Ablation studies revealed that SpliceAI2’s high performance was driven by rarely used splice sites, whose characterization was facilitated by the sequencing depth achieved through large-scale RNA-seq aggregation, and whose contribution during training was reinforced through cube-root transformation (Extended Data Fig. 2).

### Modeling tissue-specific splicing

The splicing code is interpreted differently across tissues, shaped largely by differences in the activity of *trans*-acting splicing factors^24–26^. To capture these tissue-specific effects, we developed a fine-tuning framework that takes genomic sequence together with the expression levels of 147 RNA binding proteins (RBPs) in the corresponding tissue as inputs and predicts splicing in that tissue (Fig. 2a and Extended Data Fig. 3). Unlike multitask approaches that learn an implicit latent representation for each tissue, this framework represents tissue context explicitly through a biologically meaningful tissue signature.

**Fig. 2.**
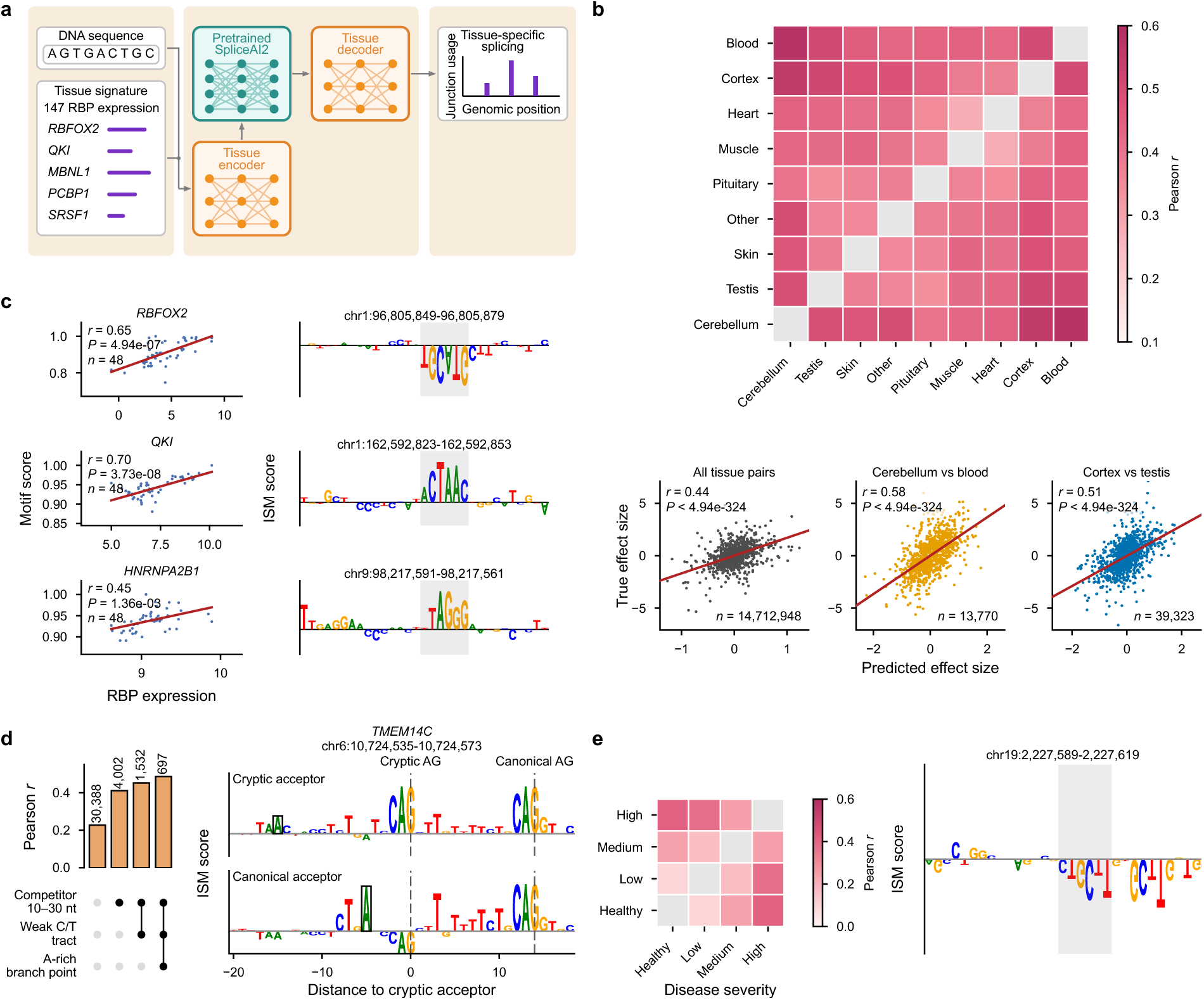
Tissue-specific splicing prediction and interpretation. **a,** Schematic of the fine-tuning framework, in which the pretrained SpliceAI2 sequence encoder is conditioned on a tissue signature comprising 147 RBP expression values through tissue encoder modules, and a tissue decoder predicts splice junction usage for the corresponding tissue. **b,** Tissue-specific splicing prediction performance, with Pearson correlations between predicted and observed junction usage differences across tissue pairs shown at the top. Aggregate and selected tissue pair comparisons are shown at the bottom, revealing stronger correlations between more distinct tissues. **c,** Interpretation of sequence features underlying tissue-specific predictions. Mean absolute ISM scores at motif sites are compared with expression of the corresponding RBPs across 48 GTEx tissues on the left, and a representative position-wise ISM profile centered on the respective motif is shown for each RBP on the right, indicating that the model uses biologically relevant features to make tissue-specific predictions. **d,** Application to tumors carrying recurrent mutations in the *SF3B1* gene. Correlations of predicted and observed junction usage differences between *SF3B1*-mutant and wild-type tumors for held-out genes, stratified by junction characteristics, are shown on the left. Correlations increased among junctions with progressively more features known to influence *SF3B1*-associated splicing. Position-wise ISM scores around competing cryptic and canonical acceptors are shown on the right, with boxes marking the branch points. **e,** Application to myotonic dystrophy disease severity. Correlations between predicted and observed junction usage differences across healthy individuals and three disease severity groups for held-out genes are shown on the left. Correlations increased with greater differences in disease severity between groups. A representative position-wise ISM profile recovering the MBNL1 motif is shown on the right.

We applied the above framework to data from 48 GTEx tissues, treating each sequence in each tissue as a distinct example during fine-tuning. We evaluated tissue-specific splicing prediction in held-out genes by comparing predicted and observed usage differences for splice junctions that were differentially used between each tissue pair, yielding 14,712,948 comparisons across all tissue pairs. Overall, SpliceAI2 predictions correlated with observed tissue-specific splicing differences (Pearson *r* = 0.44; Fig. 2b) and performed particularly well for biologically distinct tissue pairs such as cerebellum versus blood (13,770 splice junctions, Pearson *r* = 0.58) and cortex versus testis (39,323 splice junctions, Pearson *r* = 0.51). Fine-tuning the pretrained model clearly outperformed training from scratch, while conditioning on a biologically meaningful tissue signature marginally improved performance relative to one-hot encoding of tissue identity (Extended Data Fig. 4a). These results suggest that tissue-specific splicing is better captured as context-dependent modulation of a shared splicing code.

To investigate the sequence basis of SpliceAI2’s tissue-specific predictions, we performed *in silico* mutagenesis (ISM) around putative splicing factor binding sites within splice junctions and quantified the resulting changes in splicing for 48 tissues. For RBFOX, QKI, and HNRNPA2B1, these effects were strongly correlated with the expression of the corresponding splicing factors across tissues (Pearson *r* = 0.45–0.70; Fig. 2c), and the underlying sequence patterns matched the established motifs of all three factors^27–29^. Similar results were observed for additional RBPs (Extended Data Fig. 4b), demonstrating that SpliceAI2 learns biologically meaningful sequence determinants of tissue-specific splicing.

We evaluated whether SpliceAI2 could be adapted beyond healthy tissues to disease states characterized by altered spliceosome activity. We first adapted SpliceAI2 to tumors carrying recurrent mutations in the *SF3B1* gene, one of the most frequently mutated splicing factor genes in human cancer^30,31^. Following fine-tuning on RNA-seq data from 18 *SF3B1*-mutant and 62 *SF3B1* wild-type uveal melanoma tumors^32^, predicted and observed *SF3B1*-associated splicing differences were correlated in held-out genes, particularly for splice junctions characterized by competing acceptor sites 10-30 nucleotides apart, a weak polypyrimidine tract, and an A-rich branch point (Pearson *r* = 0.49; Fig. 2d)^33^. In a representative example, ISM identified distinct upstream branch points associated with the canonical and cryptic splice sites. We then adapted SpliceAI2 to myotonic dystrophy, in which loss of MBNL1 activity causes progressive splicing dysregulation^34^. Following fine-tuning on RNA-seq data from 11 healthy individuals and 44 affected individuals spanning three disease severity groups^34^, predicted and observed MBNL1-associated splicing differences were correlated in held-out genes (mean Pearson *r* = 0.26), and correlations increased between groups with greater differences in disease severity (Pearson *r* = 0.42; Fig. 2e).

We considered the challenging task of predicting tissue-specific variant effects. As observed with ABSplice^35^ and ABSplice2^36^, differences in reference splicing between tissues were highly predictive of tissue-specific variant effects, with no additional benefit from tissue-specific predictions (Extended Data Fig. 4c).

### Functional effects in human populations

We examined whether predicted changes in splicing translated to detectable effects in large human population datasets. We first analyzed WGS data from more than 627,000 individuals in the Genome Aggregation Database (gnomAD), Trans-Omics for Precision Medicine (TOPMed) program, and UK Biobank^37–39^, leveraging the depletion of splice-altering variants at common allele frequencies as a signature of purifying selection. We compared variants that were unobserved in the three cohorts with common variants (allele frequency ≥ 0.01%) after matching for trinucleotide context, with analyses stratified by variant position and probability of loss-of-function intolerance (pLI). Across all stratifications, increasing SpliceAI2 scores were associated with progressively greater depletion, approaching that of nonsense variants at high scores (Fig. 3a,b). Within high pLI genes, SpliceAI2 outperformed all other models in capturing the relationship between predicted splicing disruption and allele count (Fig. 3c).

**Fig. 3.**
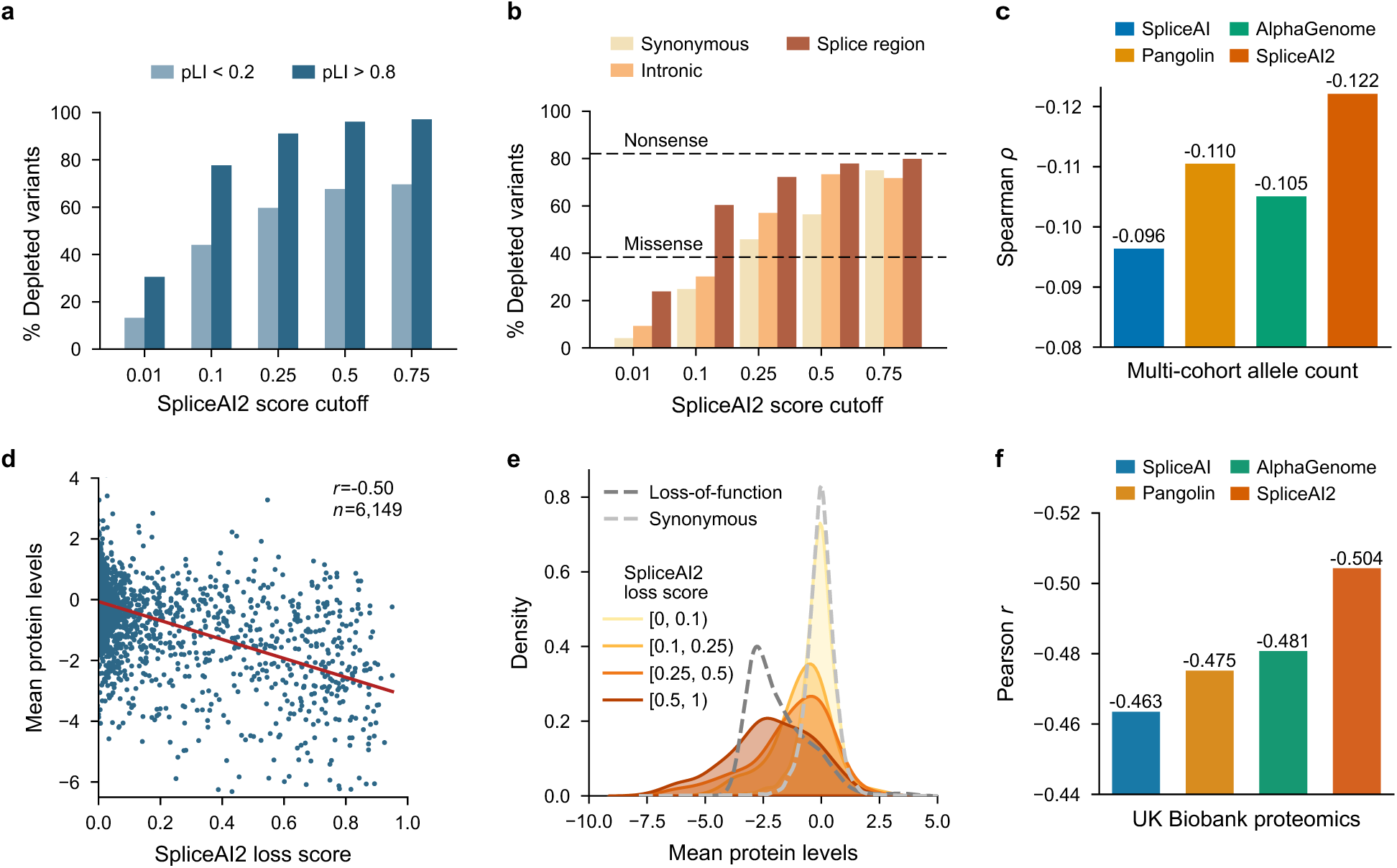
SpliceAI2 scores reflect population constraint and protein level effects. **a,b,** Percentage depletion of common variants in gnomAD, TOPMed, and UK Biobank, relative to a matched set of unobserved variants across SpliceAI2 score cutoffs, stratified by pLI and variant position, showing progressively greater depletion at higher cutoffs. Dashed lines indicate depletion levels for missense and nonsense variants. **c,** Comparison of model performance using Spearman correlation with multi-cohort allele counts in high pLI genes. More negative values indicate greater depletion of high-scoring variants. **d,** Mean carrier plasma protein levels versus SpliceAI2 loss score for rare splice region variants (allele frequency ≤ 0.1%) across 1,656 genes with plasma proteomic measurements in UK Biobank. **e,** Distribution of mean carrier plasma protein levels for rare splice region variants binned by SpliceAI2 loss score. With increasing loss score, distributions shifted from levels observed for synonymous variants toward those observed for loss-of-function variants, both indicated by dashed lines. **f,** Model comparison based on Pearson correlation with mean carrier plasma protein levels.

We then assessed whether predicted splice disruption was associated with reduced protein abundance using WGS and plasma proteomic measurements from 36,764 UK Biobank participants, restricting the analysis to splice region variants to minimize effects driven by other mechanisms. Across 6,149 rare splice region variants in 1,656 genes, SpliceAI2 loss scores were strongly negatively correlated with mean protein levels among carriers (Pearson *r* = −0.50; Fig. 3d). Stratification by SpliceAI2 loss score revealed a progressive reduction in carrier protein levels, from distributions resembling synonymous variants at low scores to those approaching loss-of-function variants at high scores (Fig. 3e). Among all models evaluated, SpliceAI2 most accurately predicted the quantitative effects of rare splice region variants on protein abundance (Fig. 3f).

### Contribution to rare genetic disease

We investigated the disease relevance of predicted splice-altering variants using WGS and matched phenotype data from the Genomics England (GEL) 100,000 Genomes Project^40^. For each proband, we used GEL PanelApp to assign disease-associated panel genes based on phenotype^41^, restricting the analysis to 7,504 probands with monoallelic disorders and fewer than 100 panel genes. We tested whether prioritized variants were enriched in panel genes relative to non-panel genes, using phenotype shuffling among probands to establish the background expectation. We quantified enrichment using the odds ratio (OR) of panel match and the number of excess panel-matched variants, both calculated by comparing prioritized variants in panel and non-panel genes between the observed and phenotype-shuffled cohorts (Fig. 4a). When calculating enrichments, we excluded missense and protein-truncating variants to isolate the contribution of cryptic splicing.

**Fig. 4.**
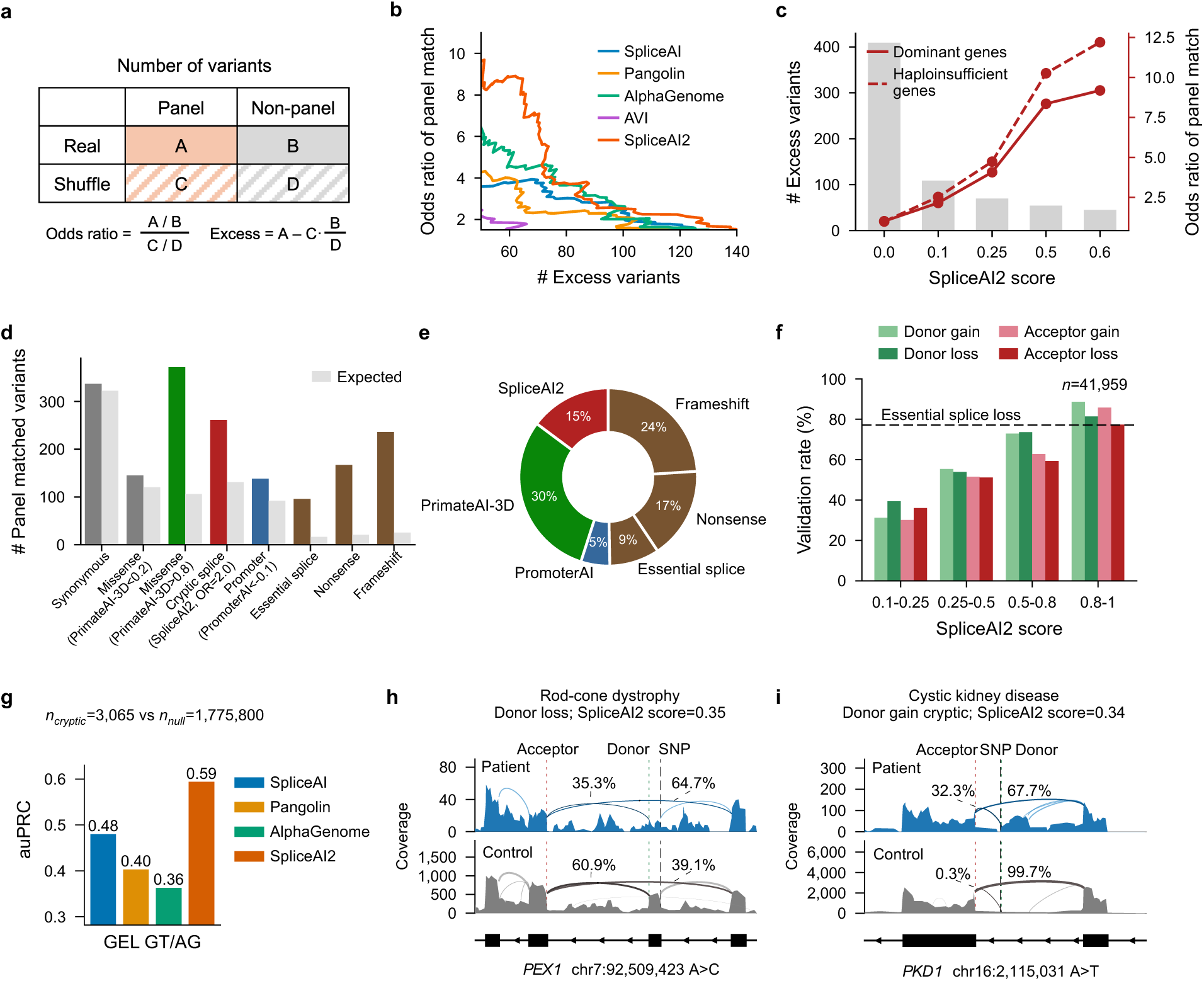
Cryptic splice variants predicted by SpliceAI2 are enriched in disease genes. **a,** Schematic of the panel match enrichment test. For a given score threshold, a 2×2 contingency table compares observed and phenotype-shuffled prioritized variants by panel match status. Enrichment is evaluated in rare disease probands from GEL without a previously prioritized TIER1 variant and quantified using odds ratios and excess variants relative to phenotype-shuffled background expectations. **b,** Panel match odds ratio versus the number of excess variants recovered across models and score thresholds. **c,** SpliceAI2 enrichment stratified by disease gene category. Enrichment increases with more stringent thresholds and is strongest for haploinsufficient genes. **d,** Number of panel-matched variants (colored bars) versus background expectation (grey bars) across variant classes. Deleterious missense variants were identified using PrimateAI-3D > 0.8, promoter variants using PromoterAI < −0.1, and cryptic splice variants using the SpliceAI2 threshold corresponding to OR = 2.0. Synonymous, essential splice, nonsense, and frameshift variants were annotated using VEP. **e,** Same as **d**, but expressed as the percentage of excess variants attributed to each category. **f,** RNA-seq validation rates of predicted donor and acceptor gain and loss variants across SpliceAI2 score bins in the GEL RNA cohort. A variant counts as validated when junction usage in carrier samples falls in the extreme tail (≤5th for loss, ≥95th for gain), matching the predicted effect direction. Validation rates increase with score, approaching the essential singleton splice loss rate at high scores (dashed line). **g,** Model comparison for cryptic splice site detection among rare variants creating a GT or AG dinucleotide. **h,i,** RNA-seq sashimi plots validate predicted splicing changes in unresolved probands. A *PEX1* variant causes partial donor loss, reducing usage of the affected junction from 60.9% in pooled controls to 35.3% in the patient. A *PKD1* variant activates a cryptic splice donor, with 32.3% usage in the patient, compared with 0.3% in pooled controls lacking the variant.

Across score thresholds for SpliceAI2 and other splicing prediction models, with the AlphaGenome AVI score included^42^, we compared the odds ratio of panel match with the number of excess panel-matched variants and found that SpliceAI2 substantially outperformed other models across the full range of thresholds (Fig. 4b). At a fixed odds ratio of 2, SpliceAI2 recovered 133 excess variants, compared with 114 for the next best model. Consistent with predicted disruption of gene function, variants prioritized by SpliceAI2 showed the highest enrichment for haploinsufficient genes (Fig. 4c).

We repeated the analysis for additional classes of genetic variation, including protein-truncating, missense, and promoter variants, enabling direct comparison of their contributions to rare disease genetic burden (Fig. 4d). Among all excess panel-matched variants, protein-truncating variants accounted for 50% of the genetic burden, with cryptic splice variants identified by SpliceAI2 contributing an additional 15% and the remainder attributable to variants identified by PrimateAI-3D^43^ and PromoterAI^44^ (Fig. 4e). Notably, half of the cryptic splice variants identified by SpliceAI2 were located more than 50 bp into introns and would therefore typically be missed by exome sequencing.

We next leveraged blood RNA-seq data from 5,435 GEL participants to validate SpliceAI2 predictions. Among 41,959 predicted splice-altering variants (SpliceAI2 > 0.1), validation rates increased progressively with SpliceAI2 score threshold and reached levels comparable to essential splice variants at high thresholds (Fig. 4f). SpliceAI2 also considerably outperformed competing models (auPRC, 0.59 versus 0.48 for SpliceAI) in predicting cryptic splice site activation by rare variants creating a GT or AG dinucleotide (Fig. 4g). When we applied the phenotype-based panel gene criteria to participants with RNA-seq data, allowing up to 500 panel genes, a SpliceAI2 threshold of 0.25 yielded 23 prioritized rare variants, of which 17 showed evidence of the predicted splicing change, including disruption of an existing splice site in the *PEX1* gene associated with rod-cone dystrophy and activation of a cryptic splice site in the *PKD1* gene associated with cystic kidney disease (Fig. 4h,i and Extended Data Fig. 5).

## Discussion

Accurate interpretation of the functional consequences of genetic variation remains a central challenge in human genetics, particularly for non-coding variation. SpliceAI2 advances non-coding variant interpretation by substantially improving the characterization of variant effects on splicing. The rare disease analysis highlighted the importance of these improvements, with variants prioritized by SpliceAI2 accounting for an estimated 15% of the excess genetic burden.

Despite this progress, several opportunities remain to improve splicing prediction. Further progress is needed to predict how splice-altering variant effects vary across tissues, cell types, and developmental contexts. Single-cell transcriptomic datasets offer a promising path to resolving such context-dependent effects, provided their sparse and biased junction coverage can be effectively addressed. Large-scale perturbation datasets targeting splicing factors in diverse cellular states may help bridge the gap between sequence-driven and context-dependent splicing prediction. The increasing availability of long-read transcriptomic datasets offers additional opportunities to characterize complex splicing events and to facilitate joint modeling of splicing and polyadenylation. Emerging datasets that measure the timing and order of splicing events may help characterize the temporal dynamics of splicing. Together, these advances should enable increasingly comprehensive prediction of how genetic variation alters splicing across biological contexts.

## Methods

### Training datasets

**Short-read RNA-seq.** We compiled bulk RNA-seq data for ten mammalian species from recount3^15^ and the NCBI Sequence Read Archive (SRA), yielding 314,745 samples and 67,493,026 splice junctions (Supplementary Table 1).

Human and mouse junctions were obtained from recount3. Projects with fewer than 100 samples were excluded. Each project was classified by Llama 3.3 (via Ollama) as bulk RNA-seq, single-cell RNA-seq, or ambiguous based on its title and abstract. Only bulk RNA-seq projects were retained, with manual review for ambiguous human projects.

For the other eight species, bulk Illumina reads were identified in SRA with the NCBI Entrez Programming Utilities and filtered to exclude non-bulk transcriptomic assays. Metadata were screened for additional evidence of single-cell, spatial, or UMI-based protocols, which were excluded. Sequencing runs with an average read length below 50 nucleotides were also removed. Reads were mapped to species-specific NCBI RefSeq genomes (Supplementary Table 1) with STAR^45^ (version 2.7.11b) using annotation-free indices. Junction counts were taken from STAR *SJ.out.tab* and summed across samples within each species.

Across all species, junctions were restricted to annotated protein-coding genes using GENCODE v48 (human), vM25 (mouse), or NCBI RefSeq (other species). We retained junctions with both ends in the same annotated protein-coding gene, at least one end at an annotated donor or acceptor site, and a per-gene maximum junction count of at least 1,000.

**Long-read RNA-seq.** We downloaded all available ENCODE long-read RNA-seq datasets^14,46,47^, comprising 189 human (GRCh38) and 141 mouse (GRCm38) samples. Transcripts were restricted to protein-coding genes with gene novelty class *Known* and transcript novelty classes *Known*, *NIC*, *NNC*, and *ISM*, and were required to contain only splice sites observed in the short-read RNA-seq datasets described above.

Transcript counts were first summed across samples, merged across isoforms with identical exon chains, and subsequently merged across isoforms sharing the same intron chain, thereby treating transcripts with identical splicing patterns as equivalent. For each intron chain, transcription start and end sites were chosen from the most abundant supporting transcript. Within each gene, transcripts whose first or last exon did not overlap the first or last exon of the most abundant transcript or were supported by only a single read were removed. Finally, only genes with at least 100 reads supporting the most abundant transcript were retained. The resulting transcript set contained 14,257 human genes and 13,491 mouse genes, represented by 593,157 and 379,729 transcripts, respectively.

### SpliceAI2 model

**Inputs.** The model takes as input a 196,608 bp genomic sequence on the sense strand of the gene, represented as a 14 × 196,608 matrix. The first four channels one-hot encode the nucleotide at each position and the remaining ten channels one-hot encode the species.

**Architecture.** The 14 × 196,608 input matrix is first projected to 256 channels by a dense layer, giving a 256 × 196,608 representation. This representation is passed sequentially through 20 MetaFormer blocks^48^, each consisting of a token mixer and a channel mixer, both with residual connections. The token mixer applies batch normalization followed by a depth-wise convolution of kernel size 5 and dilation rate that doubles every two blocks. The channel mixer applies batch normalization followed by a feed-forward network with expansion ratio 4 and rectified linear unit (ReLU) hidden layer activation. Each block is preceded by an average pooling layer and followed by linear upsampling back to full base-pair resolution, with the pooling factor doubling every four blocks.

Each of the 20 block outputs is cropped to the central 65,536 positions and reduced to two channels by separate dense layers. Their average across blocks, followed by a sigmoid activation, gives the predicted splice donor and acceptor usage per position. The 1,024 positions with the highest splice site usage, taken as the maximum of the splice donor and acceptor usage, are selected and sorted by coordinate along the sense strand. The 20 block outputs are subset to the selected 1,024 positions, then layer normalized and mapped to queries and keys with the same dimension by two dense layers, all separate per block, with rotary position embeddings (RoPE) applied using coordinates. The scaled dot product of queries and keys gives a 1,024 × 1,024 matrix per block. The lower triangle of these matrices, including the diagonal, is masked out.

Their average across blocks, followed by a sigmoid activation, gives the predicted splice junction usage per pair of positions. In total, the model has approximately 13 million trainable parameters (Fig. 1a, Extended Data Fig. 1).

**Outputs.** The model outputs a 2 × 65,536 matrix of predicted splice donor and acceptor usage across the central 65,536 positions, and a 1,024 × 1,024 matrix of predicted splice junction usage for every pair of candidate splice sites, defined as the 1,024 positions among the 65,536 positions with the highest predicted splice donor or acceptor usage.

The predicted splice donor, acceptor, and junction usages define a splice graph over the candidate splice sites. Each site is a node, ordered by coordinate along the sense strand, and each pair of sites forms a directed edge representing an intron, with the upstream and downstream sites acting as splice donor and acceptor, respectively. For efficient implementation, each donor node weight is absorbed into its outgoing edge weights, and each acceptor node weight is absorbed into its incoming edge weights, yielding an equivalent representation. Thus, each edge weight is the sum of the logits of the splice donor usage at its upstream site, the splice junction usage of the pair of sites, and the splice acceptor usage at its downstream site. Consecutive sites are additionally joined by edges of weight zero representing exon continuation. A transcript corresponds to a path through this graph, a chain of non-overlapping introns separated by exonic stretches, and its score is the sum of the weights of the edges along the path.

**Training.** Splice site and splice junction usage targets come from short-read junction counts, which for each gene are converted into splice donor or acceptor usage by summing over all junctions using each position as donor or acceptor, respectively, and dividing by the largest such count in the gene. The junction counts are converted to junction usage through division by this same value. A cube root operation is applied to all usage values. At training time, the 1,024 candidate splice sites for which splice junction usage is predicted are selected from ground truth splice site usage instead of predicted splice site usage. This stabilizes training in early epochs when predictions are arbitrary and avoids materializing the 65,536 × 65,536 junction usage matrix. The splice site and splice junction outputs are trained with binary cross entropy loss against these usage targets, restricted to positions within genes.

Transcript usage targets come from long-read transcript counts, which for each gene are converted into transcript usage through division by the gene’s total transcript count. Transcript loss is the categorical cross entropy between the observed usage distribution across transcripts and the predicted distribution calculated from the splice graph. Both distributions are defined over the set of all possible transcripts in the splice graph, denoted T, whose cardinality grows exponentially in the number of candidate splice sites. Let *w_i_*_j_ be the weight of the edge from node *i* to node *j*. The score of a transcript *T* in T is,

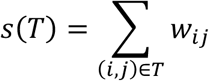

and the graph assigns it a predicted usage by normalizing over all transcripts,

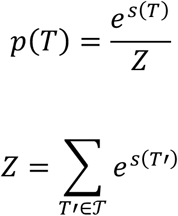

Suppose *q*(*T*) is the observed usage of transcript *T*, the categorical cross entropy between the two distributions is

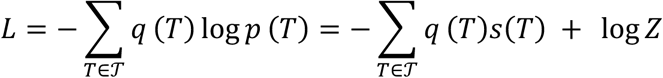

Only the observed transcripts contribute to the first term, enabling efficient computation. The second term can be computed efficiently by dynamic programming. Let *αⱼ* be the sum of the exponentiated scores of all paths through the subgraph formed by the first *j* candidate splice sites, that is,

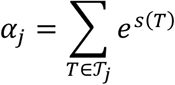

where T*ⱼ* is the set of all possible transcripts in the subgraph. Then *α*_j+1_ can be calculated as

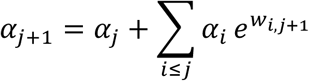

The first term corresponds to the weight zero exon continuation edge from site *j*, which extends every path of T*ⱼ* at no cost. The second term corresponds to an intron from an earlier site *i*, which extends a path of T*_i_* contributing *w_i_*_,j+1_ to its score. This gives the exact summation in quadratic time in the number of candidate splice sites.

The total loss is the sum of the splice site, splice junction, and transcript losses, with the transcript loss weighted by 10^−5^ to compensate for the large value of log *Z* and keep the different loss terms comparable in scale.

Genes on human chromosomes 21 and 22 were held out for validation, and all remaining genes were used for training. All weights were initialized from a normal distribution with a mean of zero and a standard deviation of 0.01, and all biases were initialized to zero. The model was trained for 50 epochs using the Muon optimizer for feed-forward weight parameters and AdamW for all remaining parameters^49,50^. Both optimizers used an initial learning rate of 10^−3^ and a weight decay of 0.01, with the learning rate reduced to 10^−4^ after 40 epochs. Training used bfloat16 mixed precision, a batch size of four per device across four NVIDIA H100 GPUs, gradient checkpointing, and gradient clipping at 10^−4^. We used two models trained with identical settings to form the final ensemble.

**Transcript prediction.** Transcripts are predicted by finding the highest scoring paths through the splice graph constructed from the splice donor, acceptor, and junction outputs. Viterbi decoding recovers the top scoring path exactly in quadratic time in the number of candidate splice sites.

Let *βⱼ* be the score of the highest scoring path from the first site to site *j*. Then,

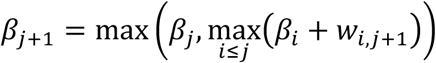

Decoding consists of a forward recursion over the sites, which computes the best score reaching each site and records the predecessor that achieved it, followed by a backtrack from the final site that reconstructs the path itself. Each path is turned into a transcript by reading its exon continuation edges as exons and the rest of its edges as introns.

Replacing the maximum in the recursion with a *k*-maximum, so that *βⱼ* holds the scores of the *k* highest scoring paths from the first site to site *j*, gives the *k* best transcripts in a single dynamic programming pass (Fig. 1b).

**Variant effect prediction.** A variant is scored by comparing predictions for the genomic sequences with reference and alternate alleles. Because the candidate splice sites used for splice junction and transcript prediction are selected from the splice site predictions, independently scoring each allele with each model in the ensemble could produce different candidate sets, particularly when the variant creates or disrupts a splice site. We therefore used a shared set of candidate splice sites for both alleles and both models in the ensemble, enabling elementwise comparison of the resulting splice junction and transcript predictions.

Both alleles are represented as 196,608 bp sense strand genomic sequences centered on the variant position and passed through the MetaFormer blocks. Splice donor and acceptor usages are predicted for both alleles and averaged over the ensemble. The candidate splice sites shared by the two alleles are the 1,024 positions whose splice site usage, maximized over splice donor and acceptor and over both alleles, is largest. Splice junction usages are then computed at these shared sites for the two alleles separately and averaged over the ensemble. This yields matched splice site and splice junction predictions for the two alleles, and variant effects are defined as the difference between the corresponding predictions for the alternate and reference alleles.

For each variant, the model reports gains and losses of splice donor, acceptor, and junction usage, retaining the ten largest effects per category. Each is reported with the predicted usage in the reference and alternate sequences and the signed distance from the variant to the affected site, or to both affected sites in the case of a junction. A single summary score is defined as the maximum predicted effect across donor gain, donor loss, acceptor gain, and acceptor loss. Junction gain and loss effects provide no net improvement and are therefore excluded from the summary score but retained for visualization.

### Evaluation of competing methods

**SpliceAI.** Variants in the GTEx and GEL GT/AG, GTEx sQTL, and UK Biobank benchmarks were scored using a modified version of SpliceAI (release 1.3.1) that reports position-resolved donor and acceptor predictions for both the reference and alternative alleles. Predictions were averaged across the five bundled models using the GRCh38 reference genome, with masking disabled. For each variant, donor or acceptor channel was selected according to the effect type, and the signed delta score (alternative minus reference) was extracted at the effect position.

Thus, positive and negative scores indicate increased and decreased predicted splice site usage, respectively. For the population genetic and GEL rare disease analyses, standard SpliceAI scores were calculated using the default settings with masking disabled.

**Pangolin.** Scores for the GTEx and GEL GT/AG, GTEx sQTL, and UK Biobank benchmarks were generated on GRCh38 with a locally modified build that left the released model weights and architecture unchanged while allowing scoring on soft-masked sequence. To obtain scores at specific splice sites, the output score cutoff was set to 0 and the per-position prediction was taken at the specified effect position. Masking was turned off so that scores remain comparable across methods. For the population genetic and GEL rare disease analyses, Pangolin was run under its default settings with masking disabled, and the reported value was the maximum absolute score across the two channels (gain and loss).

**AlphaGenome.** All AlphaGenome analyses were conducted at the University of Oxford. Predictions were generated using a 1 Mb sequence context centered on each variant. We used *score_variant()* with the recommended SPLICE_SITES, SPLICE_SITE_USAGE, and SPLICE_JUNCTIONS variant scorers to obtain variant scores and *predict_variant()* to access position-specific donor and acceptor predictions for evaluations based on specified effect positions and effect types, confirming that *score_variant()* and *predict_variant()* produced identical values for comparable outputs. Composite scores were calculated by summing the splice site and splice site usage effects with one-fifth of the splice junction effect, following the recommended scaling. We evaluated the individual splice site, splice site usage, and splice junction scores, as well as their composite, using raw and quantile outputs and maximum and mean aggregation strategies. Among 40 scoring configurations tested, the raw composite score with maximum aggregation showed the strongest average performance across the eight evaluations, although the raw splice site score with maximum aggregation performed noticeably better in GTEx GT/AG auPRC (0.63) and GEL GT/AG auPRC (0.45). We selected the raw composite score with maximum aggregation for all evaluations, as it performed best overall and is also the recommended configuration for splice variant effect prediction. AVI scores were downloaded from the AlphaGenome Atlas downloads page.

### Top transcript benchmark

We restricted the human long-read transcript set to multi-exon transcripts from genes on held-out chromosomes 21 and 22. Only genes in which the most abundant transcript had at least twice the read support of the second most abundant transcript were considered. Reconstruction was considered correct when the predicted transcript exactly matched the observed transcript from 10 bp upstream of the first splice site to 10 bp downstream of the last splice site.

Because this benchmark evaluates reference sequence predictions rather than variant effects, models were trained exclusively on human data to prevent cross-species leakage through orthologous genes. Two models were trained, one using only short-read RNA-seq data and the other using both short-and long-read RNA-seq data (Fig. 1c).

### Variant effect benchmarks

**GTEx and GEL GT/AG.** To benchmark splicing prediction models on variants that create *de novo* splice sites, we identified single nucleotide variants (SNVs) from WGS data that create a GT or AG dinucleotide, then used matched RNA-seq junction data from the same individuals as ground truth, labelling variants with junction support at the novel site as cryptic gains and those without as null. Specifically, to reduce false negatives caused by reference-mapping bias against the variant allele, novel junctions were additionally called from STAR alignments to a personalized diploid genome, in which one haplotype was the reference genome and the second haplotype carried the individual’s alternate alleles in place of the corresponding reference alleles. Novel junction support was then taken from reads aligning to the variant-carrying haplotype. We labelled variants as cryptic gains when the novel junction was observed exclusively in carriers of the variant, and, for GTEx samples, was supported by more than one sample. We defined a variant as null when it created a GT or AG dinucleotide but had no RNA-seq junction support, was carried by at least ten individuals, and fell in a gene highly expressed across all individuals, in order to ensure that the absence of a new junction reflected a genuinely inert variant rather than a lack of transcription. We applied this pipeline to GTEx and GEL samples. For GTEx samples, we also derived a quantitative usage benchmark by dividing each novel junction count by the sum of junction counts at the gene’s most abundant donor or acceptor sites.

**GTEx sQTL.** We downloaded GTEx v8 significant sQTL variant-junction pairs and corresponding SuSiE fine-mapping results for all available tissues^51,52^. Variant-junction pairs were restricted to single nucleotide variants occurring within 50 bp of either end of the associated splice junction. For each variant-junction pair, effect sizes were averaged across tissues with a significant association. Fine-mapped variants were selected by requiring PIP ≥ 0.9 in at least one tissue, whereas null variants were selected by requiring PIP < 0.1 in all tissues and an absolute mean effect size below 0.5. After excluding variants overlapping the essential splice dinucleotides, the final benchmark comprised 2,385 fine-mapped variants and 3,824 null variants (Fig. 1d).

**OpenSplice MPRA.** We evaluated 283,121 single nucleotide substitutions across 599 exons with an experimentally measured ΔPSI using SpliceAI, Pangolin, and AlphaGenome scores released with the OpenSplice dataset. To match the exon skipping predictions of the other models, we calculated the SpliceAI2 predicted effect for each variant as the average of the change in donor usage at the donor site and the change in acceptor usage at the acceptor site flanking the measured exon,

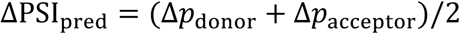

where Δ*p* = *p*_alt_ - *p*_ref_.

### Tissue-specific model architecture and fine-tuning

**Inputs.** SpliceAI2 was extended with tissue signature conditioning to predict tissue-specific splice junction usage for 48 GTEx v8 tissues (Fig. 2a). The sequence input was one-hot encoded with 1 kb of flanking sequence on each side. Unknown bases were encoded as zeros. Input was capped at 64 kb, with longer genes split into consecutive segments. The signature input was a 147-dimensional RBP expression vector, covering 79 RBPs from RBPmap^53^ and 68 from oRNAment^54^. Expression was quantified from GTEx bulk RNA-seq. Values were normalized by the trimmed mean of *M*-values (TMM) method^55^, converted to counts per million, and log_2_-transformed. The expression vector was divided by 20 and broadcast along the sequence dimension.

**Architecture.** Signature injection modules were placed before each of the 20 MetaFormer blocks to condition the model on tissue-specific information (Extended Data Fig. 3). Each injection module coupled the sequence and signature streams by bidirectional feature-wise linear modulation (bi-FiLM)^56^. A convolutional block with batch normalization and a squeeze-and-excitation gate followed. The output was scaled by a learnable weight and added to the input sequence embedding through a residual connection. A three-layer cross-attention decoder mapped the merged features to junction usage logits. Queries were formed from the sequence embeddings at anchor splice sites and added to a learned tissue embedding. The decoder used RoPE (θ = 1 × 10⁵), linear relative position biases, and query-key normalization. A bounded transformation, 10 × tanh(x/10), was applied to stabilize the output logits.

**Outputs.** For each queried anchor splice site, positions upstream of a donor anchor or downstream of an acceptor anchor were masked. A positional softmax over the remaining logits produced the distribution of tissue-specific junction usage.

**Training.** Fine-tuning minimized cross-entropy loss for tissue-specific anchor-based junction usage, weighted binary cross-entropy loss for donor and acceptor classification, and a mean squared error contrastive loss on predicted versus measured pairwise tissue differences in anchor-based junction usage. MetaFormer block weights were loaded from the pretrained multispecies SpliceAI2 model. The signature injection modules and decoder modules were randomly initialized, and the full model was then fine-tuned. Muon^49^ was used for matrices and convolutional kernels, and AdamW^50^ for biases, normalization parameters, and scalar gates. Training used a learning rate of 2 × 10⁻⁴, a weight decay of 0.01, cosine decay with warm-up, and mixed precision. The model was trained for 10 epochs on four NVIDIA H100 GPUs. Chromosomes 2, 4, 6, 8, 10–22, and X were used for training. Chromosomes 1, 3, 5, 7, and 9 were used for validation.

**Evaluation.** Predictions were averaged across two independently trained replicates, except for *in silico* mutagenesis, which used one replicate. Model performance was evaluated by Pearson correlation with two-sided *P* values on logit-transformed anchor-based junction usages, after the clipping described below.

### Tissue-specific splicing analysis

**Prediction of tissue-specific junction usage.** Ground truth was obtained from GTEx v8 short-read junction counts, summed across samples within each tissue, for protein-coding genes in GENCODE v48. Junctions were required to have a positive 75th-percentile per-sample read count, a length below 32 kb, and placement within genes. Evaluation used the held-out chromosomes 1, 3, 5, 7, and 9. For an anchor splice site with several matching sites, anchor-based junction usage was the junction count divided by the summed counts of all junctions sharing that anchor. Junctions in genes no longer than 64 kb were evaluated. For each junction, the 24 tissues of highest measured usage were paired with the 24 lowest, giving 576 tissue pairs. Junction usages were clipped to [10⁻⁶, 1 - 10⁻⁶], logit-transformed, and differenced between tissues. For each tissue pair, junctions with fewer than 1,500 anchor reads in either tissue were excluded. The heatmap pooled points across nine tissue groups, after which Pearson correlation was computed (Fig. 2b). Eight non-cortical, non-cerebellar brain regions and postmortem blood were excluded from this heatmap.

**Tissue-specific variant effect prediction.** To benchmark variants with tissue-specific splicing effects, we assembled SNVs from GTEx v8 WGS and matched multi-tissue RNA-seq (Extended Data Fig. 4c). Twenty-four tissues were aggregated into ten meta-tissues. We first restricted analysis to SNVs within 20 bp of splice sites in protein-coding genes with more than 20 supporting reads at the anchor splice site in both meta-tissues of the comparison, excluding usages of 0 or 1. We computed ΔSNV at each variant–junction pair as the carrier-minus-non-carrier difference in mean junction usage within each meta-tissue. Tissue-specific SNVs were then retained when, for at least one of the meta-tissue pairs, the absolute difference in ΔSNV between meta-tissues exceeded 0.01, at least two constituent tissues per meta-tissue showed carrier usage beyond predefined extremes, and this difference exceeded the mean ΔSNV across the pair. Reference and alternate alleles were evaluated under each tissue signature. A ridge regression (α = 1, standardized features, fivefold cross-validation grouped by chromosome) predicted variant effects from (i) measured reference usage in the two meta-tissues, or (ii) measured reference usage plus logit-transformed predicted alternate-allele probabilities in the two meta-tissues.

### RBP motifs and *in silico* mutagenesis

Six-nucleotide motifs were obtained from high-confidence human ATtRACT matrices^57^ (Fig. 2c, Extended Data Fig. 4b). For RBPs absent from ATtRACT, motifs were derived from the first six positions of the oRNAment position-weight matrix. For each RBP, the tissues of highest and lowest expression were paired for discovery.

Within each discovery pair, candidate junctions were required to differ in measured usage by more than 0.5 between the two tissues. They were also required to be shorter than 32 kb and to lie on chromosomes 1–22. Every exact motif match within a candidate junction defined a 51 bp window centered on the motif, with the motif at positions 26–31. *In silico* mutagenesis was then performed independently for each such window. The per-position ISM score was the reference value minus the mean value across the three substitutions at that position. The per-event motif score was the mean ISM score over the six motif positions. Motif strength was the motif score divided by the mean ISM score across the flanking positions located more than 10 bp from any splice site. The 400 windows with the highest motif strength per RBP were evaluated across all 48 tissues. The mean absolute per-event score per tissue was regressed on RBP expression by ordinary least squares. Expression of *RBFOX2* and *RBFOX3* was averaged, because both proteins recognize GCAUG.

### Disease-specific splicing analysis

***SF3B1*-mutant uveal melanoma (UVM).** Mutation data, gene expression, and splice junction counts from 80 UVM samples were obtained from the Genomic Data Commons (https://gdc.cancer.gov)^32^. Mutant and wild-type conditions were supplied as two 147-RBP expression vectors from bulk RNA-seq, with identical DNA input. Predicted and measured mutant-minus-wild-type anchor-based junction usage differences were compared after clipping at ε = 10⁻⁴. Only junctions with more than 500 anchor reads were retained. Competing acceptors were defined as acceptors that shared the same donor. The polypyrimidine tract metric was the C or T fraction at transcript offsets −20 to −1 from the acceptor G. The branch point metric was the adenosine fraction at offsets −18 to −8. Junctions were filtered cumulatively for a competing acceptor 10–30 nt apart, a polypyrimidine fraction below 0.65, and a branch point adenosine fraction of at least 0.2 (Fig. 2d). *In silico* mutagenesis of *TMEM14C* used the mutant signature, with per-position effects computed in log space.

**MBNL1-depleted myotonic dystrophy.** We analyzed published RNA-seq data from 55 tibialis anterior biopsies from 44 individuals with myotonic dystrophy type 1 (DM1) and 11 unaffected individuals^34^. Paired-end reads were obtained from the Sequence Read Archive (GEO GSE86356) and processed with the GTEx RNA-seq pipeline V10. Alignments to GRCh38 (GENCODE v39) were generated with STAR (version 2.7.10a) in two-pass mode. Gene-level expected counts were estimated with RSEM (version 1.3.3), then scaled by TMM^55^ and expressed as log counts per million. Splice junction usage was calculated from the number of uniquely mapping reads spanning each intron. Inferred functional MBNL levels for these biopsies were taken from Wagner et al.^58^. Tissue signatures were four 147-RBP vectors representing healthy tissue and disease of low, medium, and high severity, corresponding to progressively decreasing levels of free MBNL1. Inferred free MBNL1 expression (log2 TMM counts per million) was 9.17, 6.01, 3.67, and 1.32, respectively (*n* = 11, 15, 14, and 15). For each condition pair, junctions were required to have more than 500 supporting anchor reads and an absolute measured logit difference above 0.01. Predicted and measured junction usages were clipped at ε = 10⁻⁴ before logit transformation (Fig. 2e). *In silico* mutagenesis was applied to MBNL1-sensitive junctions that contained matching motifs.

### Population genetics analysis

We generated a benchmark of protein-coding SNVs within 30 bp of annotated splice sites comprising simulated absent variants and observed common variants (allele frequency ≥ 0.01%) from a combined cohort of gnomAD (v3.1.2), TOPMed (freeze 8), and UK Biobank (∼450k)^37–39^. Absent and common variants were trinucleotide-context matched by enforcing an approximately 1:1 class ratio within each context through resampling, and all essential splice variants were excluded.

For Fig. 3a,b, model performance was evaluated using percentage depletion based on SpliceAI2 prediction scores. Percentage depletion at each score threshold was defined as (1 - 1/OR) × 100, where OR represents the odds of variant absence among variants scoring at or above the threshold relative to those scoring below it. Depletion was stratified by gene constraint in Fig. 3a (gnomAD v4.1 pLI > 0.8 versus < 0.2) and by genomic region in Fig. 3b (synonymous, intron, and splice region variants). Missense and nonsense depletion values were computed as the odds ratio of absence relative to synonymous variants. For Fig. 3c, models were compared using Spearman correlation between per-variant prediction scores and combined multi-cohort raw allele counts.

### UK Biobank proteomics analysis

We used rare variants in a set of 1,656 protein-coding genes from the UK Biobank plasma proteomics cohort^59^. Genes were selected based on significant burden test associations between deleterious coding variants and the corresponding plasma protein (Olink) levels. For each gene, we extracted every variant within the region spanning the transcript and extending 10 kb on each side from UK Biobank WGS data (∼500,000 participants). Each variant was annotated with its consequence using the Ensembl Variant Effect Predictor (VEP)^60^ and matched to the plasma protein phenotype associated with that gene.

Plasma protein phenotypes were Olink proteomics residuals, adjusted for GWAS covariates and 90 genetic principal components (PCs), inverse rank normal transformed and refit. For each gene, we further regressed out the genotypes of coding variants with allele frequency (AF) > 0.1% (missense, protein-truncating, in-frame indel, start-or stop-altering, and canonical splice donor or acceptor consequences), and used these residuals as the phenotype. For each variant, we then identified its carriers and computed the mean and s.d. of the adjusted phenotype across carriers. Carriers of rare (AF < 0.1%) coding variants in the same gene were excluded so that non-coding signal was not confounded by co-occurring coding variation. Means and s.d. values were computed for unrelated individuals of European ancestry.

We scored every variant at the splice site nearest to the variant under the canonical annotation (GENCODE v39). The per-variant score was the model’s delta score, with loss scores given a positive sign and gain scores a negative sign. We restricted the analysis to splice region variants defined by VEP^60^, with AF ≤ 0.1% and carried by at least two individuals with measured protein levels. Essential splice variants were excluded. All model comparisons were based on the Pearson correlation between the model score and the mean protein *z*-score across carriers.

### Genomics England rare disease analysis

**Panel-matched disease gene enrichment.** We used rare variants ascertained in probands with rare disease from the Genomics England (GEL) 100,000 Genomes Project (The National Genomic Research Library, Genomics England (2024). https://doi.org/10.6084/m9.figshare.4530893; project ID 754). Each proband was associated with one or more disease gene panels (Genomics England PanelApp; accessed 25 April 2023). We restricted the cohort to the 8,778 probands whose assigned panels contained fewer than 100 genes rated with “confirmed/probable” confidence. We also excluded panel genes for which a TIER1 variant was reported in GEL v16. We retained singleton single nucleotide variants and insertions or deletions (indels) that were rare in gnomAD (minor allele frequency < 0.1%).

Variant consequences were annotated with VEP v109^60^. The analyses focused on panel genes with a dominant mode of inheritance (7,504 probands). A panel gene was classified as dominant if any of its recorded inheritance modes was monoallelic, ‘both’, X-linked dominant, imprinted, or hemizygous for genes on chromosome X in males^61^. We also compared this definition with the subset of dominant genes that are haploinsufficient^61^.

For the comparison of splice prediction models, we restricted the enrichment analysis to variants whose VEP annotated consequences were not essential splice, stop gained, start lost, frameshift, or missense. We quantified enrichment as the tendency for variants to map to genes on the proband’s disease panel (‘panel match’). To establish a null expectation, we generated ten shuffled datasets in which disease-panel assignments were permuted across probands while preserving each proband’s marginal variant set and panel size. To generate collision-minimizing shuffles, we scored a candidate permutation *π* by the energy *E*(*π*) = - ∑*_i_ J* G*S_i_*, *S_π_*_(*i*)_H, the summed Jaccard overlap between each individual’s panel gene set *S_i_* and that of the individual they are mapped to. Starting from a random permutation, we minimized *E* by Metropolis–Hastings simulated annealing, with *T*_0_ = 0.10 cooled by 0.995 per step, for 10*N* steps for *N* individuals. The lowest-energy permutation was retained. Expected null counts were taken as the mean across shuffles.

For a given model score threshold, we built a 2×2 contingency table contrasting real and shuffled data by panel match status. We computed the odds ratio OR = (A·D) / (B·C), with the terms defined in Fig. 4a. The number of excess variants recovered above the null was defined as A – C · B / D. We tested the statistical significance of the enrichment using a two-sided Fisher’s exact test. When restricting the analysis to dominant disease genes, variants in recessive genes were counted as non-panel-matched rather than discarded.

**RNA-seq validation.** To evaluate splice variant predictors, we built an RNA-seq benchmark from 5,435 probands with matched WGS and RNA-seq data. In protein-coding genes (hg38), 47,783,068 variants were called, of which we analyzed the 292,651 variants with a SpliceAI2 score > 0.1. These variants fell into two classes, each validated by a different approach. For variants creating a canonical GT donor or AG acceptor dinucleotide, we classified the variant as validated if the predicted novel junction was present in the personalized diploid-genome alignments. For the rest of the variants, we quantified the anchor usage of the affected junction (defined in the “Prediction of tissue-specific junction usage” section) and built a cohort reference distribution from samples with ≥ 10 junction reads at that anchor. A junction qualified only if ≥ 1,000 samples met this coverage requirement; splice-loss junctions additionally required non-zero baseline usage in ≥ 90% of qualifying samples. After filtering, 41,959 variant–gene pairs retained a qualifying junction. For each variant-gene pair, we computed the percentile of every carrier against the cohort reference distribution and averaged these percentiles at the most strongly affected junction; a variant was validated when the mean carrier percentile was ≤ 5th percentile for splice loss or ≥ 95th percentile for splice gain.

## Acknowledgements

The Genotype-Tissue Expression (GTEx) Project was supported by the Common Fund of the Office of the Director of the National Institutes of Health (commonfund.nih.gov/GTEx). Additional funds were provided by the NCI, NHGRI, NHLBI, NIDA, NIMH, and NINDS. Donors were enrolled at Biospecimen Source Sites funded by NCI\Leidos Biomedical Research, Inc. subcontracts to the National Disease Research Interchange (10XS170), Roswell Park Cancer Institute (10XS171), and Science Care, Inc. (X10S172). The Laboratory, Data Analysis, and Coordinating Center (LDACC) was funded through a contract (HHSN268201000029C) to The Broad Institute, Inc. Biorepository operations were funded through a Leidos Biomedical Research, Inc. subcontract to Van Andel Research Institute (10ST1035). Additional data repository and project management were provided by Leidos Biomedical Research, Inc. (HHSN261200800001E). The Brain Bank was supported by supplements to University of Miami grant DA006227. Statistical Methods development grants were made to the University of Geneva (MH090941 & MH101814), the University of Chicago (MH090951, MH090937, MH101825, & MH101820), the University of North Carolina - Chapel Hill (MH090936), North Carolina State University (MH101819), Harvard University (MH090948), Stanford University (MH101782), Washington University (MH101810), and to the University of Pennsylvania (MH101822). The datasets used for the analyses described in this manuscript were obtained from dbGaP at http://www.ncbi.nlm.nih.gov/gap through dbGaP accession number phs000424.v9.p2.

We gratefully acknowledge the participants of the National Genomic Research Library (NGRL), whose contributions made this research possible. Secure access to the NGRL under project ID 754 was provided by Genomics England, which delivers the NGRL in partnership with NHS England, and is wholly owned by the UK Department of Health and Social Care. The NGRL contains participants’ health data collected by the NHS as part of their care, along with samples and data from their participation in research, for which fully informed consent has been obtained. This includes genomic and clinical data provided through the NHS Genomic Medicine Service, as well as data obtained through research studies, including the 100,000 Genomes Project and the Generation Study, both of which are delivered in partnership with the NHS, and from other research cohorts involving external collaborators.

The Wellcome Trust, Cancer Research UK and the Medical Research Council have also funded research infrastructure. This research has been conducted using the UK Biobank (Resource Application Number 33751).

## Author contributions

K.J. conceived the study, led model development, and supervised the project. J.C. prepared all training and benchmark datasets. X.L. developed and evaluated tissue-specific and disease-specific models. Y.Z. led the UK Biobank and Genomics England analyses. G.C., G.P., P.P.F., N.W., A.B., J.F.M., and Y.C. assisted with these analyses. I.I.T., A.J.R., and K.C. contributed to model development and implementation. Y.K. and E.M.W. carried out AlphaGenome analyses. S.J.S. supervised the AlphaGenome analyses. R.M. provided project oversight. K.K.F. conceived and supervised the project.

## Data and code availability

Source code, pretrained model weights, and precomputed variant effect predictions within protein-coding genes for all possible single nucleotide variants (4 billion) and for indels observed in human populations (150 million) are available at https://github.com/Illumina/SpliceAI2.

Data from the National Genomic Research Library (NGRL) used in this research are available within the secure Genomics England Research Environment. Access to NGRL data is restricted to adhere to consent requirements and protect participant privacy. Data used in this research include: variants from /genomes/by_date/ or extracted from aggV2 variant calls (/gel_data_resources/main_programme/aggregation/aggregate_gVCF_strelka/aggV2/genomic_d ata/), RNA-seq raw fastq files, mapped bam files, gene level expression quant.gene.sf files, and junction counts SJ.out.tab files from /gel_data_resources/RNASeq_data/Rare_Disease/, and patient disease status and panel gene information from PanelApp.

Access to NGRL data is provided to approved researchers who are members of the Genomics England Research Network, subject to institutional access agreements and research project approval under participant-led governance. For more information on data access, visit: https://www.genomicsengland.co.uk/research

## Extended data figures

**Extended Data Fig. 1.**
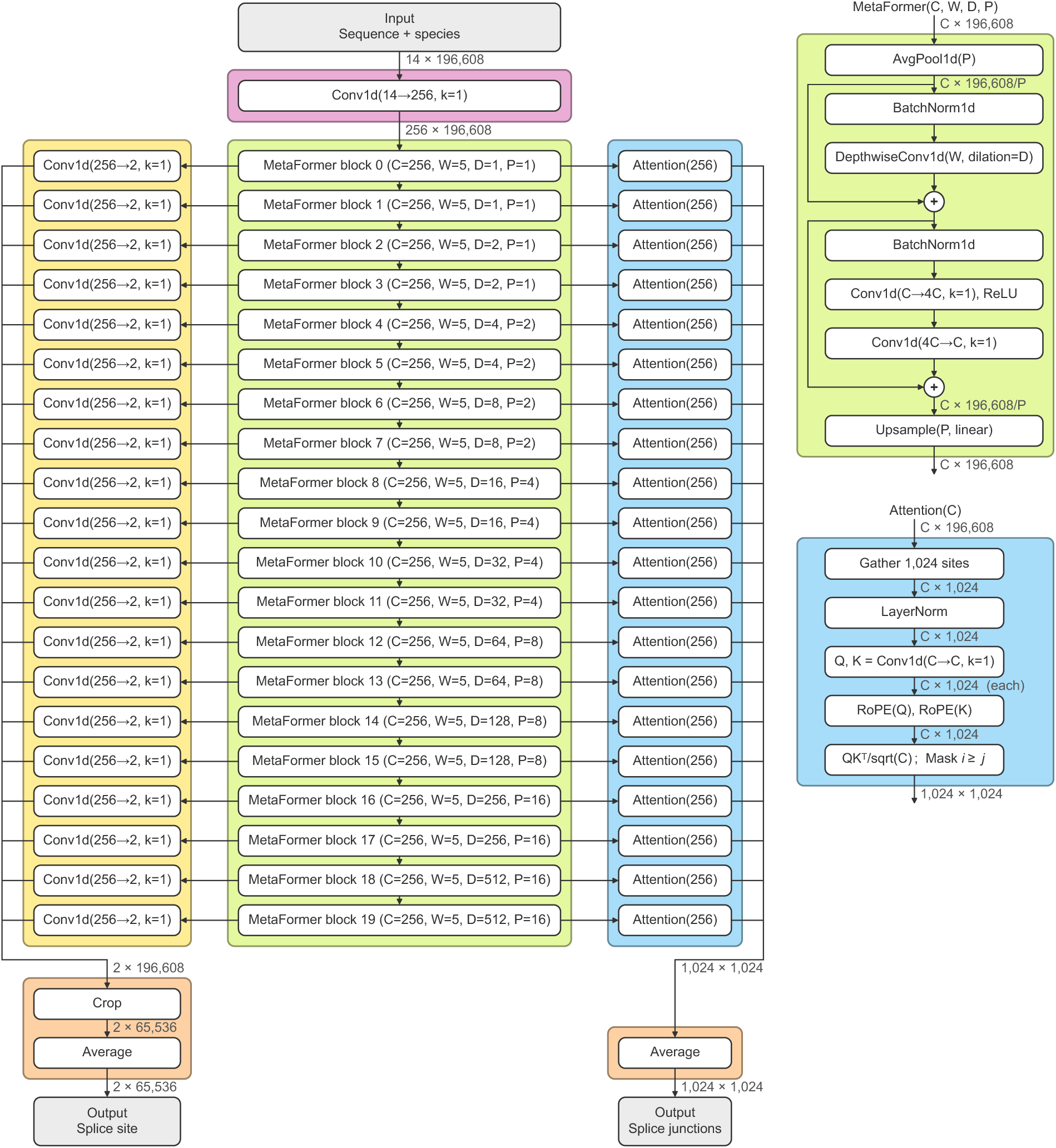
SpliceAI2 model architecture. The model processes 196,608 bp of genomic sequence represented by 14 channels, comprising four one-hot nucleotide channels and ten one-hot species channels. The input is embedded into 256 channels (pink) and processed by 20 MetaFormer blocks (green) with increasing pooling and dilation rates. Splice site heads (yellow) predict splice site logits across the central 65,536 bp, and splice junction heads (blue) predict pairwise splice junction logits among the 1,024 highest scoring splice site positions. The final splice site and splice junction outputs are obtained by averaging the predictions across the 20 blocks, followed by a sigmoid activation.

**Extended Data Fig. 2.**
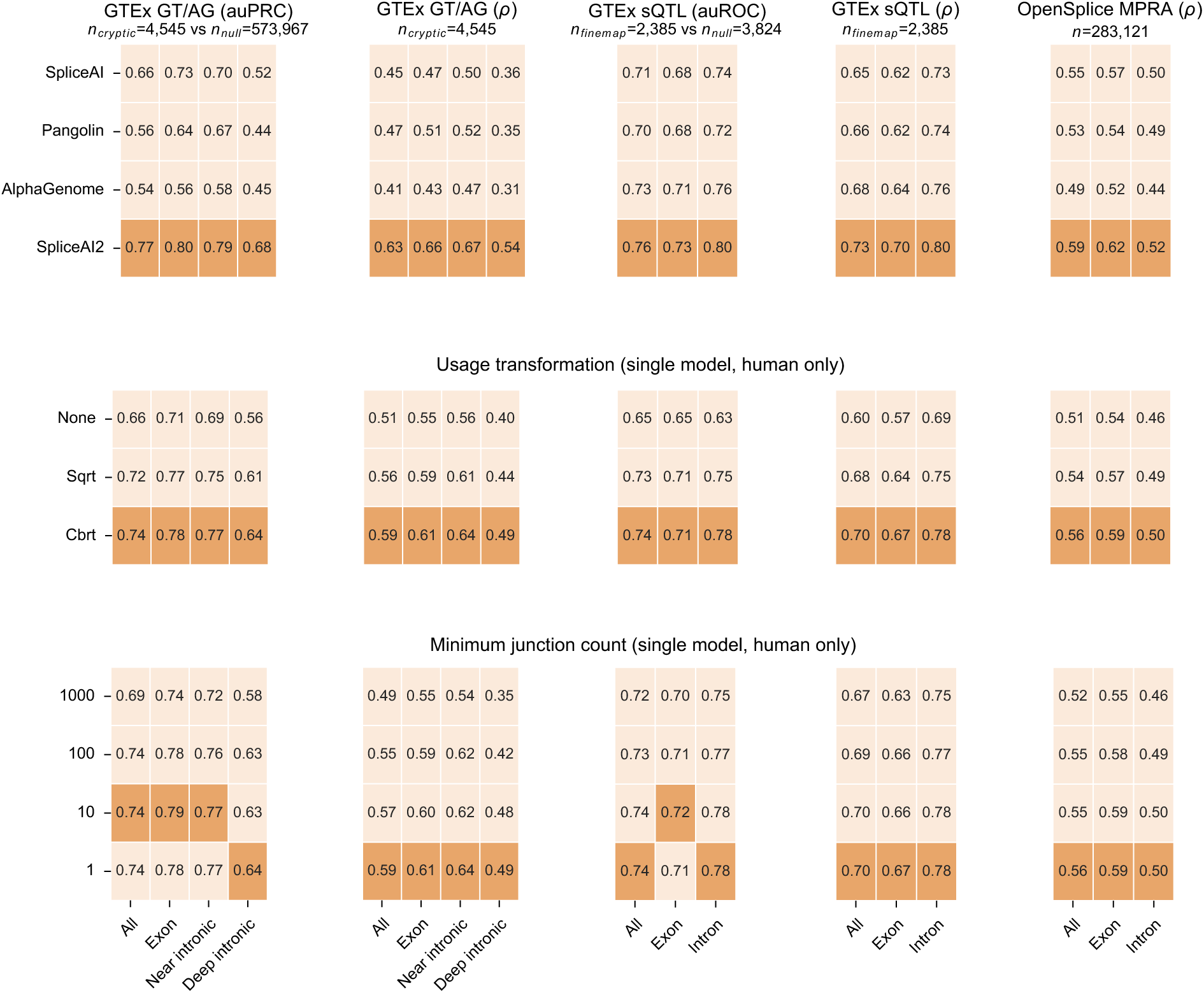
Model performance comparison and ablation studies. Performance on the variant effect benchmarks in Fig. 1d is stratified by genomic region, including exonic, near intronic, and deep intronic (≥500 bp from nearest exon). Ablation studies demonstrate the contribution of usage transformation and junction count filtering.

**Extended Data Fig. 3.**
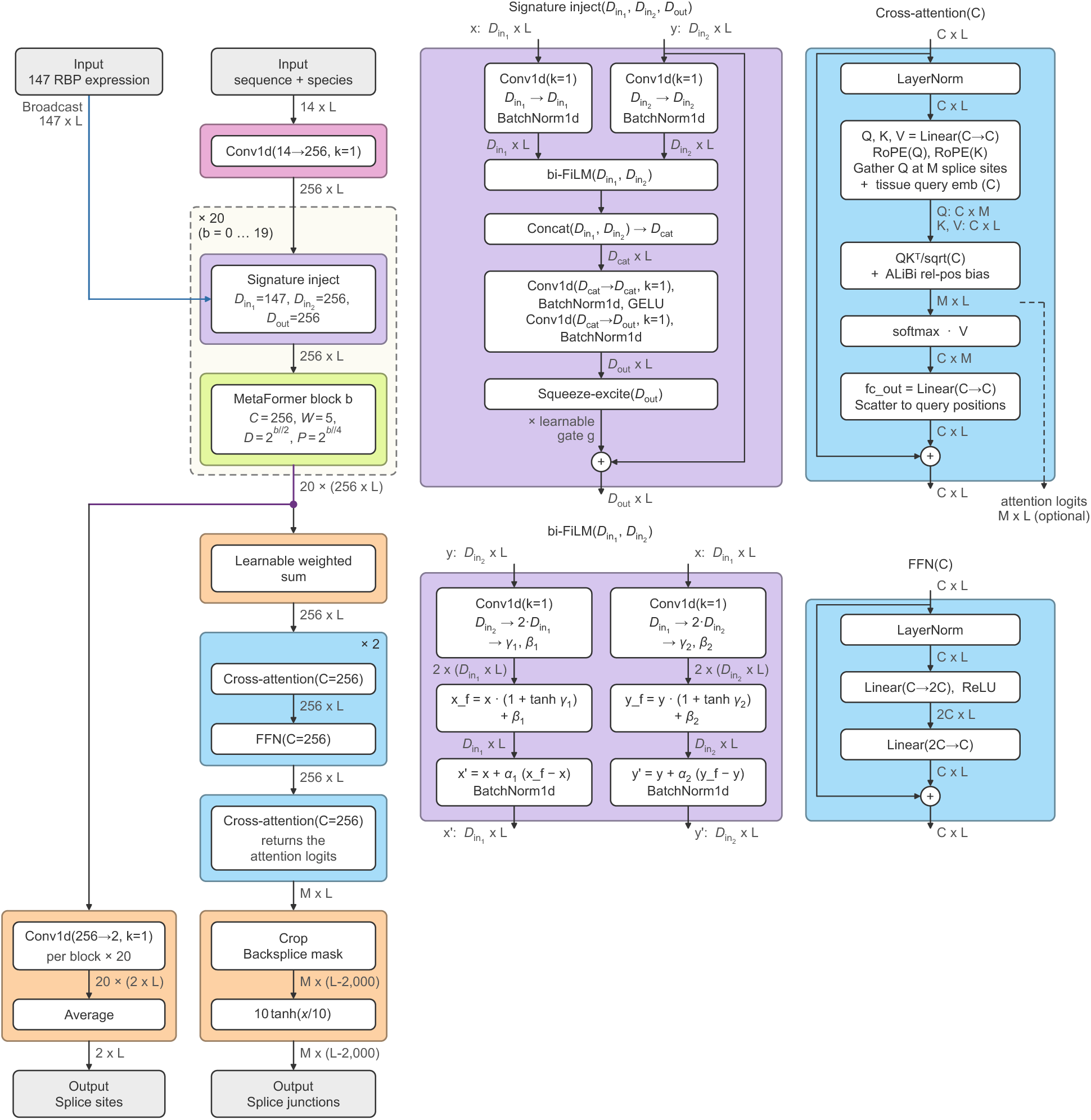
Tissue-specific SpliceAI2 model architecture. In addition to genomic sequence and species identity, the tissue-specific model receives as input a tissue signature comprising 147 RBP expression values. The sequence representation is conditioned on the tissue signature through signature injection modules (purple) before each MetaFormer block (green), whose outputs are merged by a learnable weighted sum and passed through a three-layer decoder (blue) to generate tissue-specific splice junction outputs.

**Extended Data Fig. 4.**
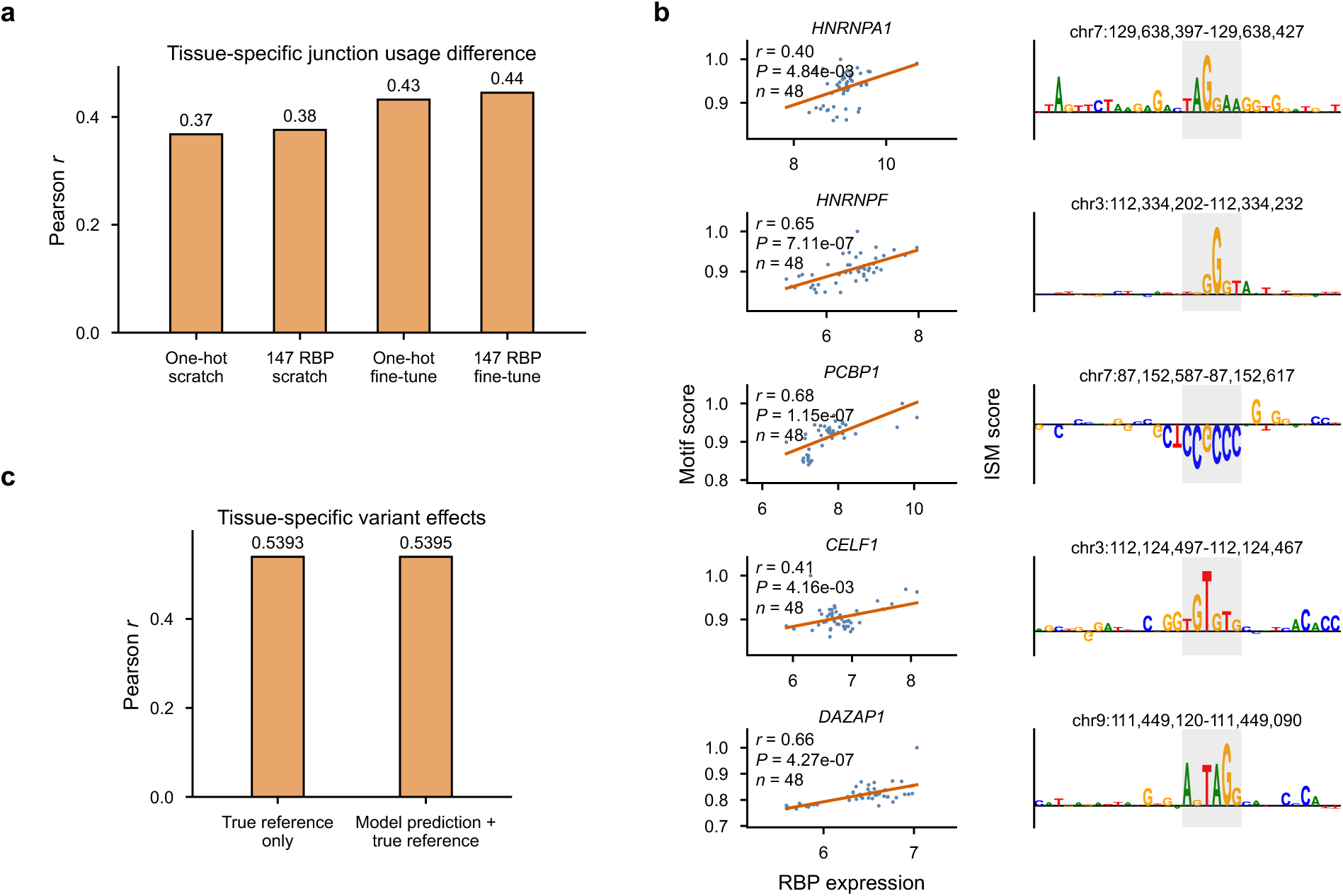
Tissue-specific splicing prediction analyses. **a,** Tissue-specific junction usage difference prediction performance across training strategies, highlighting improved performance with fine-tuning and tissue signatures. **b,** Same analysis as in Fig. 2c, shown for five additional RBPs. **c,** Tissue-specific variant effect prediction performance. Pearson correlation between observed effects and those estimated by ridge regression using reference splice junction usage probabilities alone or together with model predictions.

**Extended Data Fig. 5.**
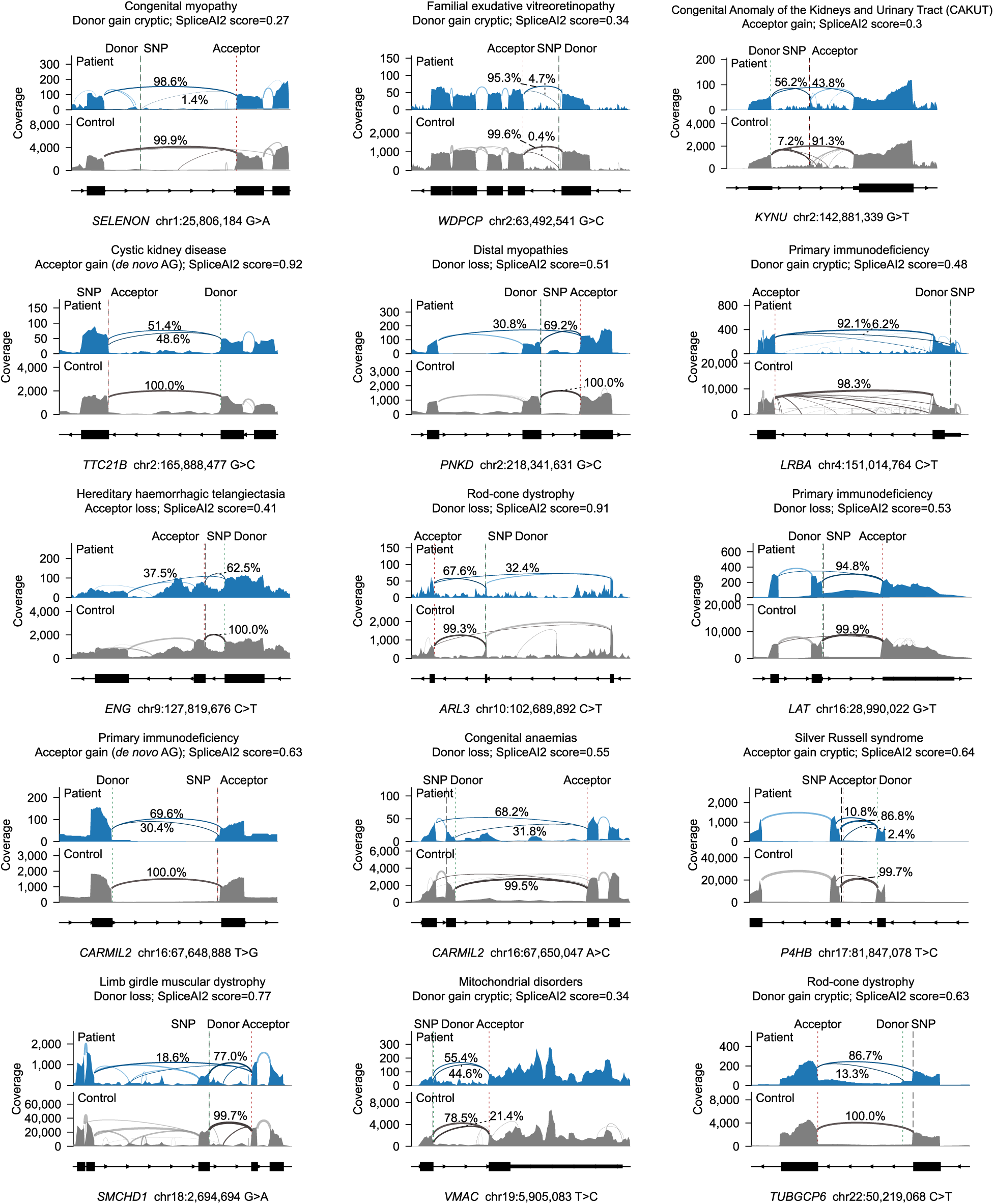
RNA-seq validated splice-altering variants from unresolved GEL probands. Sashimi plots are shown for unresolved probands with monoallelic disorders and fewer than 500 panel genes, each carrying a rare variant with a SpliceAI2 score > 0.25 in one of these genes. Each proband is compared with 20 pooled controls lacking the variant.

## References

1 Wang, Z. & Burge, C. B. Splicing regulation: from a parts list of regulatory elements to an integrated splicing code. RNA 14, 802–813 (2008). 10.1261/rna.876308

2 Barash, Y. et al. Deciphering the splicing code. Nature 465, 53–59 (2010). 10.1038/nature09000

3 Cooper, T. A., Wan, L. & Dreyfuss, G. RNA and disease. Cell 136, 777–793 (2009). 10.1016/j.cell.2009.02.011

4 Scotti, M. M. & Swanson, M. S. RNA mis-splicing in disease. Nat Rev Genet 17, 19–32 (2016). 10.1038/nrg.2015.3

5 Li, Y. I. et al. RNA splicing is a primary link between genetic variation and disease. Science 352, 600–604 (2016). 10.1126/science.aad9417

6 100000 Genomes Project Pilot Investigators et al. 100,000 Genomes Pilot on Rare-Disease Diagnosis in Health Care - Preliminary Report. N Engl J Med 385, 1868–1880 (2021). 10.1056/NEJMoa2035790

7 Ellingford, J. M. et al. Recommendations for clinical interpretation of variants found in non-coding regions of the genome. Genome Med 14, 73 (2022). 10.1186/s13073-022-01073-3

8 Jaganathan, K. et al. Predicting Splicing from Primary Sequence with Deep Learning. Cell 176, 535–548 e524 (2019). 10.1016/j.cell.2018.12.015

9 Walker, L. C. et al. Application of the Acmg/Amp Framework to Capture Evidence Relevant to Predicted and Observed Impact on Splicing: Recommendations from the Clingen Svi Splicing Subgroup. medRxiv (2023). 10.1101/2023.02.24.23286431

10 Zeng, T. & Li, Y. I. Predicting RNA splicing from DNA sequence using Pangolin. Genome Biol 23, 103 (2022). 10.1186/s13059-022-02664-4

11 Xu, C. et al. Reference-informed prediction of alternative splicing and splicing-altering mutations from sequences. Genome Res 34, 1052–1065 (2024). 10.1101/gr.279044.124

12 Avsec, Z. et al. Advancing regulatory variant effect prediction with AlphaGenome. Nature 649, 1206–1218 (2026). 10.1038/s41586-025-10014-0

13 Chao, K. H., Mao, A., Salzberg, S. L. & Pertea, M. Splam: a deep-learning-based splice site predictor that improves spliced alignments. Genome Biol 25, 243 (2024). 10.1186/s13059-024-03379-4

14 Encode Project Consortium. An integrated encyclopedia of DNA elements in the human genome. Nature 489, 57–74 (2012). 10.1038/nature11247

15 Wilks, C. et al. recount3: summaries and queries for large-scale RNA-seq expression and splicing. Genome Biol 22, 323 (2021). 10.1186/s13059-021-02533-6

16 Sarawagi, S. & Cohen, W. W. Semi-Markov Conditional Random Fields for Information Extraction. Advances in Neural Information Processing Systems 17 (2004).

17 Burge, C. & Karlin, S. Prediction of complete gene structures in human genomic DNA. J Mol Biol 268, 78–94 (1997). 10.1006/jmbi.1997.0951

18 Gelfand, M. S. & Roytberg, M. A. Prediction of the exon-intron structure by a dynamic programming approach. Biosystems 30, 173–182 (1993). 10.1016/0303-2647(93)90069-o

19 Majoros, W. H., Lebeck, N., Ohler, U. & Li, S. Improved transcript isoform discovery using ORF graphs. Bioinformatics 30, 1958–1964 (2014). 10.1093/bioinformatics/btu160

20 Gabriel, L., Becker, F., Hoff, K. J. & Stanke, M. Tiberius: end-to-end deep learning with an HMM for gene prediction. Bioinformatics 40 (2024). 10.1093/bioinformatics/btae685

21 Holst, F. et al. Helixer: ab initio prediction of primary eukaryotic gene models combining deep learning and a hidden Markov model. Nat Methods 23, 732–739 (2026). 10.1038/s41592-025-02939-1

22 The GTEx Consortium et al. The GTEx Consortium atlas of genetic regulatory effects across human tissues. Science 369, 1318–1330 (2020). doi:10.1126/science.aaz1776

23 Quarantani, G. et al. OpenSplice: the impact of half a million mutations on the alternative splicing of 600 human exons. bioRxiv (2026). 10.64898/2026.05.22.727141

24 Fu, X. D. & Ares, M., Jr. Context-dependent control of alternative splicing by RNA-binding proteins. Nat Rev Genet 15, 689–701 (2014). 10.1038/nrg3778

25 Ule, J. & Blencowe, B. J. Alternative Splicing Regulatory Networks: Functions, Mechanisms, and Evolution. Mol Cell 76, 329–345 (2019). 10.1016/j.molcel.2019.09.017

26 Tao, Y., Zhang, Q., Wang, H., Yang, X. & Mu, H. Alternative splicing and related RNA binding proteins in human health and disease. Signal Transduct Target Ther 9, 26 (2024). 10.1038/s41392-024-01734-2

27 Martinez, F. J. et al. Protein-RNA Networks Regulated by Normal and ALS-Associated Mutant HNRNPA2B1 in the Nervous System. Neuron 92, 780–795 (2016). 10.1016/j.neuron.2016.09.050

28 Galarneau, A. & Richard, S. Target RNA motif and target mRNAs of the Quaking STAR protein. Nat Struct Mol Biol 12, 691–698 (2005). 10.1038/nsmb963

29 Jbara, A. et al. RBFOX2 modulates a metastatic signature of alternative splicing in pancreatic cancer. Nature 617, 147–153 (2023). 10.1038/s41586-023-05820-3

30 Liu, Z. et al. Mutations in the RNA Splicing Factor SF3B1 Promote Tumorigenesis through MYC Stabilization. Cancer Discov 10, 806–821 (2020). 10.1158/2159-8290.CD-19-1330

31 Seiler, M. et al. Somatic Mutational Landscape of Splicing Factor Genes and Their Functional Consequences across 33 Cancer Types. Cell Rep 23, 282–296 e284 (2018). 10.1016/j.celrep.2018.01.088

32 Robertson, A. G. et al. Integrative Analysis Identifies Four Molecular and Clinical Subsets in Uveal Melanoma. Cancer Cell 33, 151 (2018). 10.1016/j.ccell.2017.12.013

33 Darman, R. B. et al. Cancer-Associated SF3B1 Hotspot Mutations Induce Cryptic 3’ Splice Site Selection through Use of a Different Branch Point. Cell Rep 13, 1033–1045 (2015). 10.1016/j.celrep.2015.09.053

34 Wang, E. T. et al. Transcriptome alterations in myotonic dystrophy skeletal muscle and heart. Hum Mol Genet 28, 1312–1321 (2019). 10.1093/hmg/ddy432

35 Wagner, N. et al. Aberrant splicing prediction across human tissues. Nat Genet 55, 861–870 (2023). 10.1038/s41588-023-01373-3

36 Wagner, N. et al. Aberrant splicing prediction during human organ development. bioRxiv (2025). 10.1101/2025.07.16.665183

37 Chen, S. et al. A genomic mutational constraint map using variation in 76,156 human genomes. Nature 625, 92–100 (2024). 10.1038/s41586-023-06045-0

38 Taliun, D. et al. Sequencing of 53,831 diverse genomes from the NHLBI TOPMed Program. Nature 590, 290–299 (2021). 10.1038/s41586-021-03205-y

39 U. K. Biobank Whole-Genome Sequencing Consortium. Whole-genome sequencing of 490,640 UK Biobank participants. Nature 645, 692–701 (2025). 10.1038/s41586-025-09272-9

40 Genomics England. The National Genomic Research Library v5.1, <https://www.genomicsengland.co.uk/> (2020).

41 Martin, A. R. et al. PanelApp crowdsources expert knowledge to establish consensus diagnostic gene panels. Nat Genet 51, 1560–1565 (2019). 10.1038/s41588-019-0528-2

42 Cheng, J. et al. AlphaGenome Atlas: in silico mutagenesis of the entire human genome improves prioritization and interpretation of non-coding variants. (2026). https://storage.googleapis.com/deepmind-media/DeepMind.com/Blog/alphagenome-atlas-a-predictive-map-of-every-possible-dna-letter-change-in-the-human-genome/alphagenome-atlas.pdf

43 Gao, H. et al. The landscape of tolerated genetic variation in humans and primates. Science 380, eabn8153 (2023). 10.1126/science.abn8197

44 Jaganathan, K. et al. Predicting expression-altering promoter mutations with deep learning. Science 389, eads7373 (2025). 10.1126/science.ads7373

45 Dobin, A. et al. STAR: ultrafast universal RNA-seq aligner. Bioinformatics 29, 15–21 (2013). 10.1093/bioinformatics/bts635

46 Kagda, M. S. et al. Data navigation on the ENCODE portal. Nat Commun 16, 9592 (2025). 10.1038/s41467-025-64343-9

47 Luo, Y. et al. New developments on the Encyclopedia of DNA Elements (ENCODE) data portal. Nucleic Acids Res 48, D882–D889 (2020). 10.1093/nar/gkz1062

48 Yu, W. et al. MetaFormer is Actually What You Need for Vision. 2022 IEEE/CVF Conference on Computer Vision and Pattern Recognition (CVPR), 10809–10819 (2022). 10.1109/CVPR52688.2022.01055

49 Liu, J. et al. Muon is Scalable for LLM Training. arXiv (2025). 10.48550/arXiv.2502.16982

50 Loshchilov, I. & Hutter, F. Decoupled Weight Decay Regularization. arXiv preprint arXiv:1711.05101 (2017). 10.48550/arXiv.1711.05101

51 Wang, G., Sarkar, A., Carbonetto, P. & Stephens, M. A simple new approach to variable selection in regression, with application to genetic fine mapping. J R Stat Soc Series B Stat Methodol 82, 1273–1300 (2020). 10.1111/rssb.12388

52 Zou, Y., Carbonetto, P., Wang, G. & Stephens, M. Fine-mapping from summary data with the “Sum of Single Effects” model. PLoS Genet 18, e1010299 (2022). 10.1371/journal.pgen.1010299

53 Paz, I., Kosti, I., Ares, M., Jr., Cline, M. & Mandel-Gutfreund, Y. RBPmap: a web server for mapping binding sites of RNA-binding proteins. Nucleic Acids Res 42, W361–367 (2014). 10.1093/nar/gku406

54 Benoit Bouvrette, L. P., Bovaird, S., Blanchette, M. & Lecuyer, E. oRNAment: a database of putative RNA binding protein target sites in the transcriptomes of model species. Nucleic Acids Res 48, D166–D173 (2020). 10.1093/nar/gkz986

55 Robinson, M. D. & Oshlack, A. A scaling normalization method for differential expression analysis of RNA-seq data. Genome Biol 11, R25 (2010). 10.1186/gb-2010-11-3-r25

56 Perez, E., Strub, F., de Vries, H., Dumoulin, V. & Courville, A. FiLM: Visual Reasoning with a General Conditioning Layer. Proceedings of the AAAI Conference on Artificial Intelligence 32, 3942–3951 (2018). 10.1609/aaai.v32i1.11671

57 Giudice, G., Sanchez-Cabo, F., Torroja, C. & Lara-Pezzi, E. ATtRACT-a database of RNA-binding proteins and associated motifs. Database (Oxford) 2016 (2016). 10.1093/database/baw035

58 Wagner, S. D. et al. Dose-Dependent Regulation of Alternative Splicing by MBNL Proteins Reveals Biomarkers for Myotonic Dystrophy. PLoS Genet 12, e1006316 (2016). 10.1371/journal.pgen.1006316

59 Sun, B. B. et al. Plasma proteomic associations with genetics and health in the UK Biobank. Nature 622, 329–338 (2023). 10.1038/s41586-023-06592-6

60 McLaren, W. et al. The Ensembl Variant Effect Predictor. Genome Biol 17, 122 (2016). 10.1186/s13059-016-0974-4

61 Sevim Bayrak, C., et al. Identification of discriminative gene-level and protein-level features associated with pathogenic gain-of-function and loss-of-function variants. Am J Hum Genet 108, 2301–2318 (2021). 10.1016/j.ajhg.2021.10.007

